# The mitochondrial RNA extrusion-induced innate immunity is regulated by N6-methyladenosine machinery

**DOI:** 10.64898/2026.08.27.747681

**Authors:** Qing Hu, Yifei Zhang, Shixuan Wang, Xueping Zeng, Fengyu Wang, Ting Fu, Yueqing Chen, Jianxin Lyu, Mijia Lu

## Abstract

Mitochondrial RNA (mtRNA) released into the cytosol functions as a damage associated molecular pattern that activates pattern-recognition receptor (PRR)-mediated inflammation, yet its release mechanisms and cytoplasmic fate remain poorly understood. Here we report that chemical Abt-373-treatment and Vesicular stomatitis virus (VSV) infection induce mtRNA extrusion through Bax/Bak and VDAC1 channels, accompanied by mtDNA release. Extruded mtRNA in A549 cells activates multiple cytosolic PRRs, including RIG-I, MDA5, TLR3/7/8, and PKR, each contributing differentially to the innate immune signaling. Analysis of GEO datasets and methylated RNA immunoprecipitation (MeRIP) assays further reveals that mtRNA carries methyladenosine (m^6^A) modification. m^6^A machinery proteins are involved in the cytoplasmic retention time of mtRNA and its interaction with RIG-I, thereby modulating mtRNA-induced innate immunity. Thus, our work establishes *in vitro* models of mtRNA extrusion, and highlights m^6^A-dependent modulation as a potential therapeutic target for mtRNA-driven inflammation.

## INTRODUCTION

Mitochondria are ubiquitous double-membrane-bound organelles in eukaryotic cells. Beyond their fundamental role in energy production, they are integral to multiple biological processes, including intracellular signal pathways that regulates apoptosis, calcium homeostasis, and innate immune responses. Within the mitochondrial matrix, the organelle harbors its own circular double-stranded DNA genome (mtDNA) and a dedicated set of transcriptional machinery. Mitochondrial transcription employs a non-canonical initiating nucleotide and generates long polycistronic RNA precursors spanning the entire genome^1^. These precursors are subsequently processed within mitochondrial RNA granules (MRGs) by the RNase P complex and RNase Z, yielding 37 transcripts including 2 rRNAs, 22 tRNAs, and 13 mitochondrial mRNAs (mt-mRNAs). mtRNA degradation is mediated by the degradosome, which consists of the SUV3 like RNA helicase and polynucleotide phosphorylase (PNPase, encoded by PNPT1), localized in D-foci.

Mitochondrial dysfunction can lead to aberrant transcription or replication, as well as mtDNA damage. Such abnormal mtDNA can be extruded from mitochondria into the cytosol through the voltage-dependent anion channel (VDAC1)^2^ or the Bax/Bak macropores on the outer mitochondrial membrane (OMM)^3^. Although mtDNA is a self-derived molecule, its cytosolic presence renders it a damage-associated molecular pattern (DAMP), which can be recognized by pattern recognition receptors (PRRs), such as cyclic GMP-AMP synthase (cGAS) in cytosol and Toll-like receptor 9 (TLR9) in endosome, activating innate immune response characterized by type I interferon (IFN) production^4–6^. Persistent activation of cGAS-STING and LTR9-MyD88 pathways lead to autologous cell damage, manifesting in autoimmune or chronic inflammatory diseases such as type II diabetes, heart failure^7^, Huntington’s disease^8^, and hepatocellular carcinoma^9^. In addition, infections by pathogens such as Herpes Simplex Virus (HSV), SARS-CoV-2, Measles virus (MeV), Modified Vaccinia Ankara (MVA), and Respiratory Syncytial Virus (RSV) have been reported to induce mtDNA release through distinct mechanisms^10–14^.

The extrusion of double-stranded mtRNA (ds-mtRNA) is also implicated in chronic inflammatory conditions. Defects in the mtRNA degradation machinery, including SUV3 and PNPase, or poly(A) polymerase (MTPAP) and the the RNA stability factor LRPPRC, have been experimentally shown to promote mRNA accumulation and release^15,16^. Cytosolic mtRNA can be recognized by RNA-sensing PRRs, including protein kinase R (PKR)^17–19^, members of retinoic acid-inducible gene I (RIG-I) like receptors (RLRs) family such as melanoma differentiation-associated protein 5 (MDA5) and RIG-I^15,20,21^, and endosomal TLR3^20^, thereby activating multiple innate immune signaling pathways. Notably, several RNA sensor, such as RIG-I and TLR3/7/8, are sensitive to N6-methyladenosine (m^6^A) modification: m^6^A-modified exogenous RNAs (e.g., synthetic or viral RNA) often fail to robustly activate these PRRs-mediated responses^22–24^. Thus m^6^A has been recognized as an evolutionarily adapted mechanism by which viruses evade innate immunity. Multiple methylation modifications such as m^5^C, m^2^2G, m^6^2A, and m^1^A have been identified on mtRNA and are involved in regulating the precursor polycistronic transcripts’ processing, maturation, degradation and cytosolic release^25–29^. However, direct evidence for m^6^A modification on mammalian mtRNA remains limited until recently. In 2024, a study demonstrated that the methyltransferase METTL17-catalyzed m^6^A modification on human mt-mRNAs^30^. This finding prompted us to investigate whether mtRNA is broadly modified by m^6^A in mitochondria and whether such modification influences the ability of extruded mtRNA to evade recognition by PRRs.

In this study, besides the chemicals- and genes deficiency-resulted mtRNA extrusion models, we demonstrate that viral infection also promotes mtRNA release, contributing to an additional layer of innate immune activation that restricts viral replication. Combining experimental data with analysis of publicly available datasets, we show that mtRNA undergose m^6^A modified, like what nuclear-encoded transcripts do. Furthermore, this modification plays a notable role in facilitating the evasion of cytosolic mtRNA from innate immune recognition.

These findings highlights the potential of targeting m6A machinery as a therapeutic strategy for mtRNA extrusion–associated autoimmunity and related pathological conditions.

## RESULTS

### ABT-737 induced mtRNA extrusion together with mtDNA through Bax/Bak channel

Chemical drug mixture containing Abt-737, S63845 and Q-VD-Oph (Abt-S-Q) was used by McArthur on HeLa cells to induce mtDNA extrusion^3^. Abt-737 is a potent Bcl-2, Bcl-xL and Bcl-w inhibitor, S63845 is a potent and selective myeloid cell leukemia 1 (MCL1) inhibitor, and Q-VD-Oph is an irreversible pan-caspase inhibitor with potent antiapoptotic properties. Here we used the mixture (Abt-S-Q) to treat A549 cells for 0, 20, 40 and 60 min, and under the confocal microscope the cells were visualized with antibodies specific for outer mitochondrial membrane (OMM) protein TOMM20 (green) and dsRNA (J2, red), which indicates the mtRNA, and DAPI for nuclear staining. The red spots presented since 20 min post treatment and as time expanded the red signal strengthened and tended to escape the envelop of OMM (TOMM20, green). At 60 min post treatment partial dsRNA was free of OMM envelop and reached the area of cytosol (Figure 1A). As a control, the mtDNA-extrusion was also observed since 20 min post Abt-S-Q-treatment. Here the dsRNA-specific antibody J2 was replaced with the one recognizing TFAM (green), which is reported to be permanently packed with mtDNA^31^ and was observed to leave the wrap of TOMM20 (red) (Figure 1B). At 0, 20, 40, and 60 min post Abt-S-Q-treatment increased mtRNA (Figure 1C) and mtDNA (Figure 1D) were detected from the mitochondria-excluded cytoplasm portion (Figure 1E). All the 13 mitochondrial transcripts and the mtDNA increased at 8 h post treatment, and then dramatically dropped at 16 h in the mitochondria-excluded A549 cytoplasm (*S.*Figure 1A-C), indicating the released mtRNA and mtDNA undergoes degradation in cytoplasm or even export out of cells. In A549 cells Abt-S-Q-treatment induced higher transcription of type I and type III interferons (IFN-β and IFN-λ, respectively), pro-inflammatory cytokines (IL-6 and TFN-α), and ADAR1p150, which was derived from IFN-induced alternative splicing and demonstrating type I IFN secretion (Figure 1F). Immunoblotting revealed the enhanced ADAR1p150 expression together with enhanced phosphorylated interferon regulatory factor 3(pIRF3), which is involved in multiple PRRs-initiated signaling pathways (Figure 1G), suggesting that the chemical drug-induced mitochondrial nucleic acids (mtNAs) release activated cytosolic PRRs and resulted in innate immunity activation. As another cell line THP-1 was treated with Abt-S-Q, similar type I and III IFN responses were activated (*S*. Figure 1D), and in mitochondria-free cytoplasm a fluctuation of mtRNA and mtDNA was observed during the 16 h post treatment (*S.* Figure 1E-G). To eliminate the mtDNA-induced innate immune response, the purified mtRNA was treated with DNase and then transfected into A549 at the dose of 0, 10 and 100 ng/well. A dose-dependent innate immunity, including pIRF3 and the transcription of IFN-β and IL-6, was observed at 12 h post transfection (Figure 1H, I).

**Figure 1.**
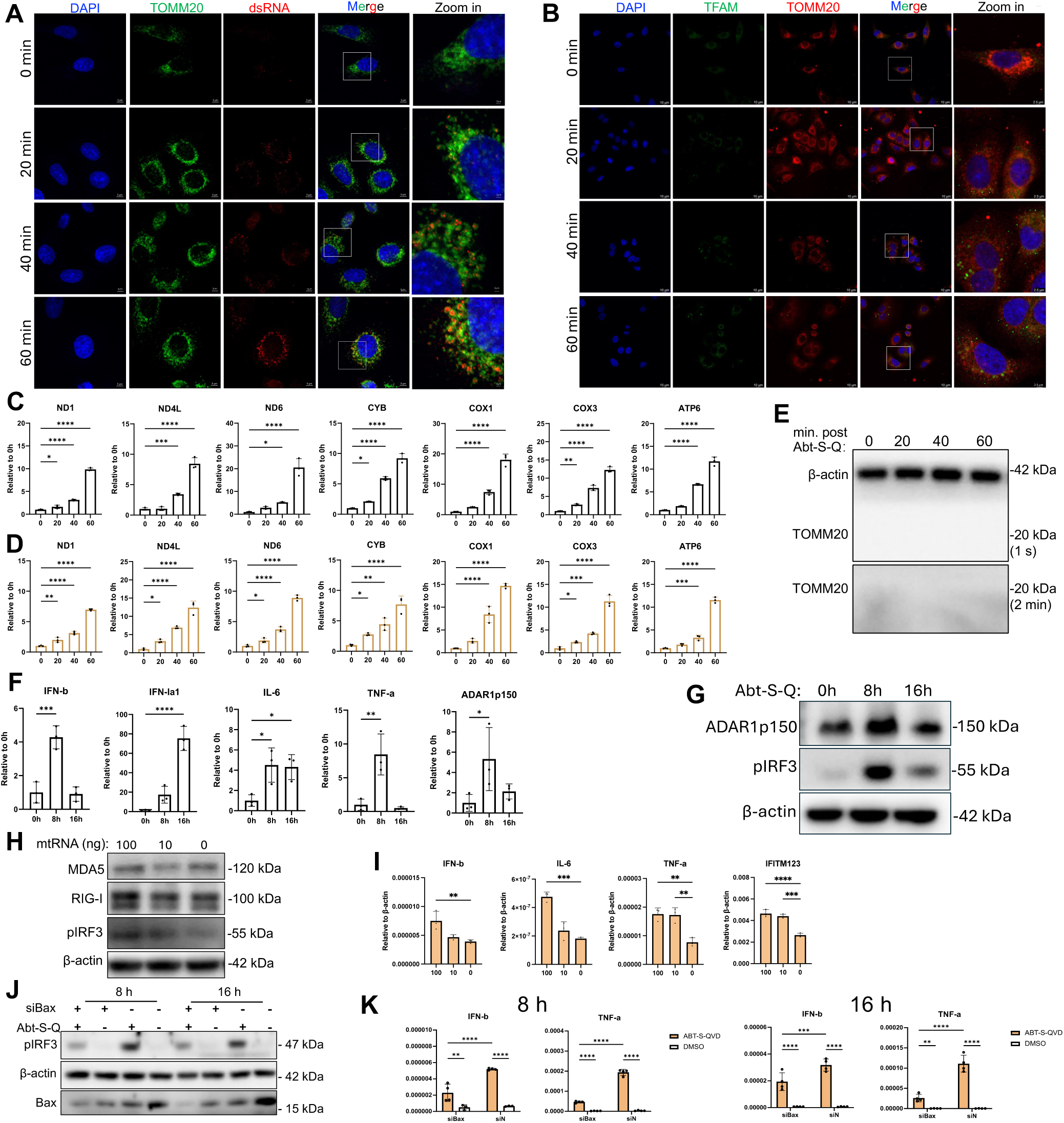
The chemical treatment induces mtRNA extrusion. (A and B) A549 cells were treated with mixture of chemical drugs (Abt-S-Q) containing Abt-737 (5 μM), S63845 (25 μM) and Q-VD-Oph (20 μM) at 37°C for 0, 20, 40 and 60 min followed by immunofluorescence assay (IFA) for mitochondrial double strand RNA (mt-dsRNA) (A) and mtDNA (B) extrusion detection. (C-E) At 0, 20, 40 and 60 min, the Abt-S-Q-treated A549 cells were subject to the collection of mitochondria-free cytoplasm for mtRNA (C) and mtDNA (D) quantification by RT-qPCR and qPCR, respectively, and the mitochondria-exclusion was testified by immunoblotting of TOMM20 (E). (F and G) A549 cells were treated with Abt-S-Q for 0, 8 and 16 h. The total RNA was extracted for the transcription quantification of IFN-β, IFN-λ, IL-6, TFN-α, and IFN-induced ADAR1p150 with RT-qPCR (F), and the cell lysate was used for phosphorylate IRF3 (S386, pIRF3) and ADAR1p150 detection with the internal reference β-actin by immunoblotting (G). (H and I) The purified mtRNA was transfected into A549 cells, 12 h later the activation of innate immunity was tested by immunoblotting (H) and RT-qPCR (I). (J and K) In A549, siRNA-introduced Bax-knockdown and Abt-S-Q-induced pIRF3 was determined by immunoblotting (J), the transcription of IFN-β and TFN-α was evaluated by RT-qPCR (K) at 8 and 16 h. Data presented as mean ± SD. n = 3 biological replicates. One-way ANOVA and two-way ANOVA were used. \**p* < 0.05, ** *p* < 0.01, *** *p* < 0.001, **** *p* < 0.0001.

When Bax was knockdown by siRNA-transfection, the Abt-S-Q-treatment resulted less mtRNA and mtDNA release (*S.* Figure 1H, I), milder pIRF3 and decreased transcription of IFN-β and TNF-α compared with siN control (Figure 1J and K). Thus the activated Bax/Bak pore is the main channel to export the mtNAs in the context of Abt-S-Q-treatment.

### VSV-infection induced mtRNA extrusion through MAVS induced mitochondrial integrity disruption

It was reported that mtDNA entered cytoplasm during virus infection through multiple mechanisms, such as mitochondria-targeting viral protein resulted mitochondrial network disruption and enhanced membrane permeability^13,14^. Vesicular stomatitis virus (VSV), a nonsegmented negative-strand (NNS) RNA virus, is capable of infecting numerous cell lines, reproducing abundantly in the cytoplasm, and finally resulting in host cell apoptosis and necrosis. In our study, a GFP-expressing recombinant VSV (rVSV-GFP) was applied to infect A549, and the viral propagation was evaluated by GFP-imaging and the viral glycoprotein (G) expression (Figure 2A and B). The infection induced dramatic innate immunity characterized by enhanced expression of pIRF3 and RIG-I, and transcription of IFN-β and cytokines, as well as type I IFN-induced IFITMs, due to the NNS RNA virus genomic RNA itself is perfect ligand of RIG-I (Figure 2B and C)^23^. Using mitoTracker, the OMM-specific fluorescent dye, mitochondrial fragmentation was observed at 8 h post infection (HPI) (Figure 2D). In the mitochondria-excluded cytosolic portion, copies of mtRNA and mtDNA were found to increase significantly over time until 16 h compared to 0 h (Figure 2E and F), indicating that VSV infection resulted in mitochondrial network fragmentation. Similar phenomena have been reported during infections with HSV-1, Measles virus, Dengue virus and Influenza A virus (IAV), although these studies primarily focused on mtDNA extrusion^10,13,14,32^. It was notable that rVSV-GFP infection resulted in dramatic downregulation of MAVS in membrane-associated fractions at the late stage (16 HPI) (Figure 2B and G), without a corresponding decrease in MAVS transcription (Figure 2H), but with more cleaved molecules (55 kDa) in the cytoplasm (Figure 2G). This suggests that MAVS undergoes a degradation, likely as a mechanism to restrain excessive innate immune response. MAVS degradation is accompanied by hexokinase 2 (HK2) loss from either membrane or cytoplasmic protein portions (Figure 2G). The OMM-located HK2 contributes greatly to mitochondrial homeostasis, like inhibiting excessive reactive oxygen species (ROS) accumulation, sustaining mitochondrial membrane potential (MMP), and restraining abnormal activation of VDAC1 by formation of MAVS-HK2-VDAC1 heterotrimer^33–37^. Upon VSV-infection, cellular MMP decreased and mitochondrial ROS (mtROS) increased markedly (Figure 2I and J), supporting that the virus leads to loss of OMM proteins and mitochondrial disruption through multiple mechanisms, thereby allowing mtRNA to enter the cytoplasm. During VSV-infection, VDAC1 knockdown reduced the release of both mtRNA and mtDNA (Figure 2K), resulting in a modest attenuation of innate immune responses (Figure 2L) and a slightly recovery of VSV replication (Figure 2M). Similarly, Bax knockdown partially inhibited mtNAs extrusion and the subsequent innate immunity, while enhancing viral replication (Figure 2N-P). These findings suggest that released mtNAs contribute to infection-induced innate immunity through both RNA sensor- and DNA sensor-mediated signaling pathways. On the contrary, when the innate immunity was established by mtRNA-extrusion in the SUV3- and PNPT1-knockdown cells (*S.* Figure 2A-C), the subsequent VSV infection was dramatically restrained (*S.* Figure 2C and D).

**Figure 2.**
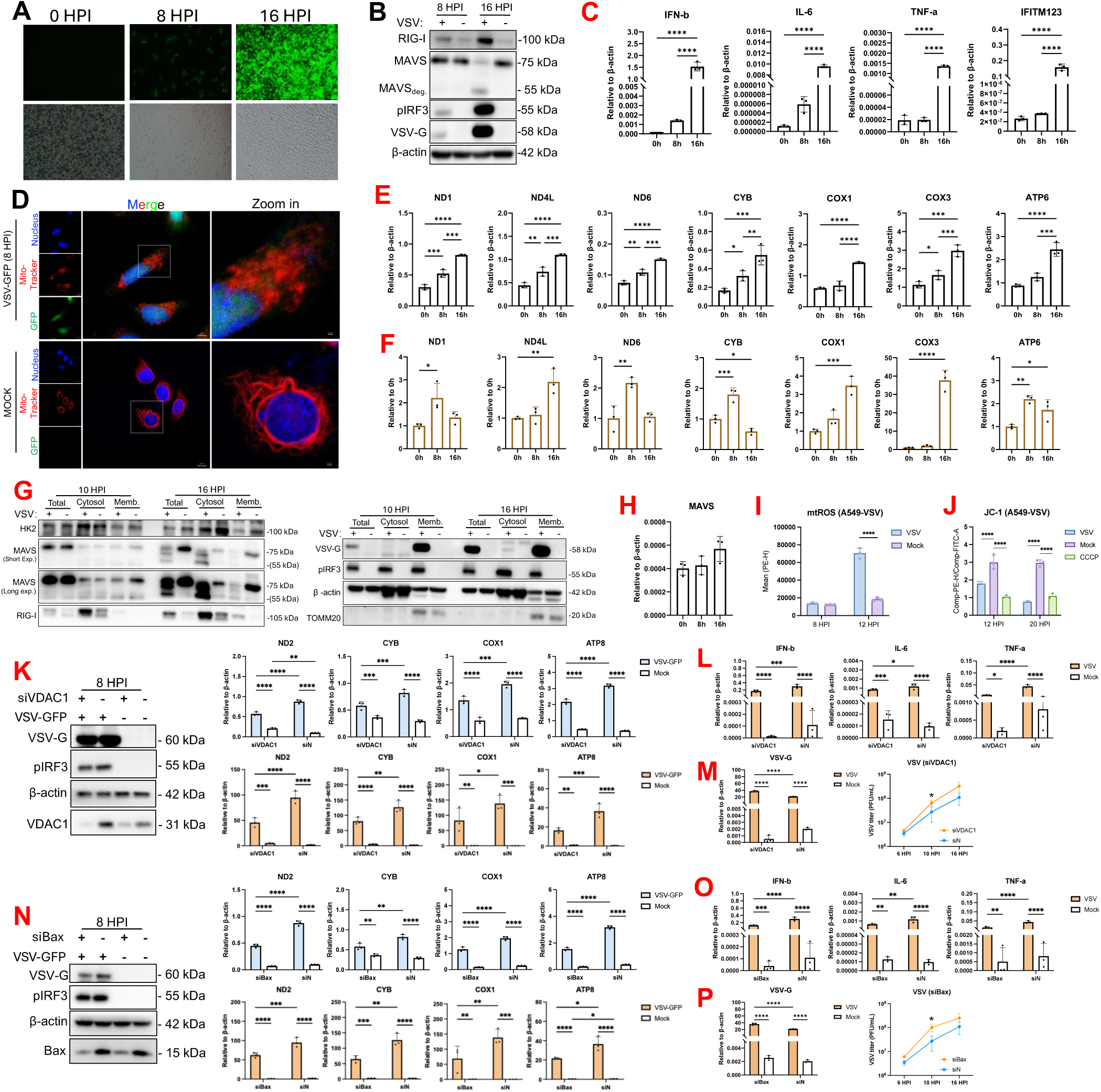
VSV-infection induces mtRNA extrusion. (A) A549 cells were infected with recombinant VSV-GFP (rVSV-GFP) and the cells were observed and imaged under fluorescent microscop at 0, 8 and 16 h post infection (HPI). (B) A549 cells were infected by rVSV-GFP at a multiple of infection (MOI) of 0.005 and then lysed at 8 and 16 HPI. The lysate was used for the indicated target proteins detection by immunoblotting. (C) A549 cells were infected with VSV with a MOI of 0.005, the total RNA was collected at 0, 8 and 16 HPI for quantification of IFN-β, IL-6, TNF-α, and IFITMs by RT-qPCR. (D) rVSV-GFP-infected (MOI= 0.01) or mock-infected A549 cells were incubated with MitoTracker and Hoechst 33342 for mitochondria and nucleus staining, respectively, at 8 HPI. (E and F) A549 cells were infected with rVSV-GFP (MOI= 0.01) for 0, 8 and 16 h. The mtRNA (E) and mtDNA (F) in the mitochondria-excluded cytoplasm were quantified and normalized with internal reference β-actin. (G) A549 cells infected with rVSV-GFP (MOI = 0.005) or mock-infected were fractionated to obtain cytosolic and membrane-associated proteins, along with total protein lysates, for immunoblot analysis. (H) MAVS mRNA levels were quantified by RT–qPCR in total RNA from VSV-infected A549 cells (MOI = 0.005) at 0, 8, and 16 HPI. (I and J) rVSV-GFP-infected (MOI= 0.01) and mock-infected A549 cells were harvested by trypsin-digestion at the indicated time points, stained with MitoSOX Red for mtROS (I) and JC-1 for MMP (J). The data were acquired by flowcytometry and analyzed with software FlowJo. (K-M) In VDAC1-knockdown A549 cells, the released mtRNA (top panels) and mtDNA (bottom panels) were determined in mitochondria-excluded cytoplasm (K). The innate immune response was evaluated by RT-qPCR (L), and the virus replication was determined by RT-qPCR and plaque assay (M). (N-P) In Bax-knockdown A549 cells, the released mtRNA (top panels) and mtDNA (bottom panels) were determined in mitochondria-excluded cytoplasm (N), the innate immunity (O) and the virus replication (P) was evaluated. Data presented as mean ± SD. n = 3 biological replicates. Two-tailed Student’s t test, one-way ANOVA and two-way ANOVA were used. \**p* < 0.05, ** *p* < 0.01, *** *p* < 0.001, **** *p* < 0.0001.

### Extruded mtRNA is recognized by multiple PRRs molecules in cytoplasm

To distinguish which cytoplasmic PRR molecule recognizes extruded mtRNA and contributes to the subsequent innate immunity, an immunofluorescence assay (IFA) was performed at 1.5 h post Abt-S-Q-treatment and discovered predominant colocalization between released mtRNA and the cytosolic dsRNA sensors RIG-I, the colocalization is relatively poor between mtRNA and MDA5/ TLR3, and even undetectable for mtRNA-PKR (Figure 3A). In A549 siRNA-introduced PRRs knockdown resulted in different degrees of inhibition on Abt-S-Q-induced IFN and inflammatory cytokines transcription. RIG-I-knockdown resulted in less pIRF3 (Figure 3B) and decreased transcription of IFN-β, IL-6 and TNF-α for 486-, 4.2- and 113-fold, respectively, compared with siN as early as 8 h (Figure 3B). However, knockdown of another RLR family member MDA5 didn’t reduce pIRF3 and only inhibited IFN-β at early infection stage (5.9-fold drop, compared with siN), though IL-6 production was significantly inhibited at both time points (5.1- and 8.2-fold drop, compared with siN, Figure 3C). TLR3-knockdown resulted dramatic IFN-β, IL-6 and TNF-α reduction at either 8 h (29.1-, 2.3- and 15.9-fold drop) or 12 h (20.9-, 4.4- and 9.7-fold drop, Figure 3d). When MyD88, the downstream adaptor of TLR7/8, was knockdown, there’s only 1.5-fold of IFN-β decrease at 8 h, but at 12 h IFN-β, IL-6 and TNF-α presented significant drop (5.2-, 10.8- and 4.5-fold, Figure 3E). The knockdown of PKR, which was reported to capable of recognizing mtRNA^17–19^, also downregulated all three transcripts obviously at both 8 h (194.2-, 4.4- and 139-fold drop) and 12 h (Figure 3F), though the colocalization was not detected at as early as 1.5 h post treatment. As expected, when TLR4 was used as control for all the other PRRs, it didn’t affect mtRNA-induced innate immunity (Figure 3G), because TLR4 locates on cell membrane and is predominantly activated by lipopolysaccharides (LPS). Besides the transcription of innate immunity-related genes, knockdown of the PRRs correspondently downregulated the IFN-β secretion in the culture medium (Figure 3H). The further RIP assay confirmed that the overexpressed RIG-I, MDA5, PKR, and TLR3/7/8, but not TLR4, can physically bind the extruded mtRNA upon Abt-S-Q treatment with different efficiency (Figure 3I). Thus our results prove that not only RLRs, but also TLRs and PKR are involved in the mtRNA recognition, activating innate immunity through multiple signaling pathways. Therefore, knockdown of any single PRR is insufficient to fully abrogate innate immune activation, due to compensatory signaling by other RNA sensors as well as contributions from mtDNA-activated pathways.

**Figure 3.**
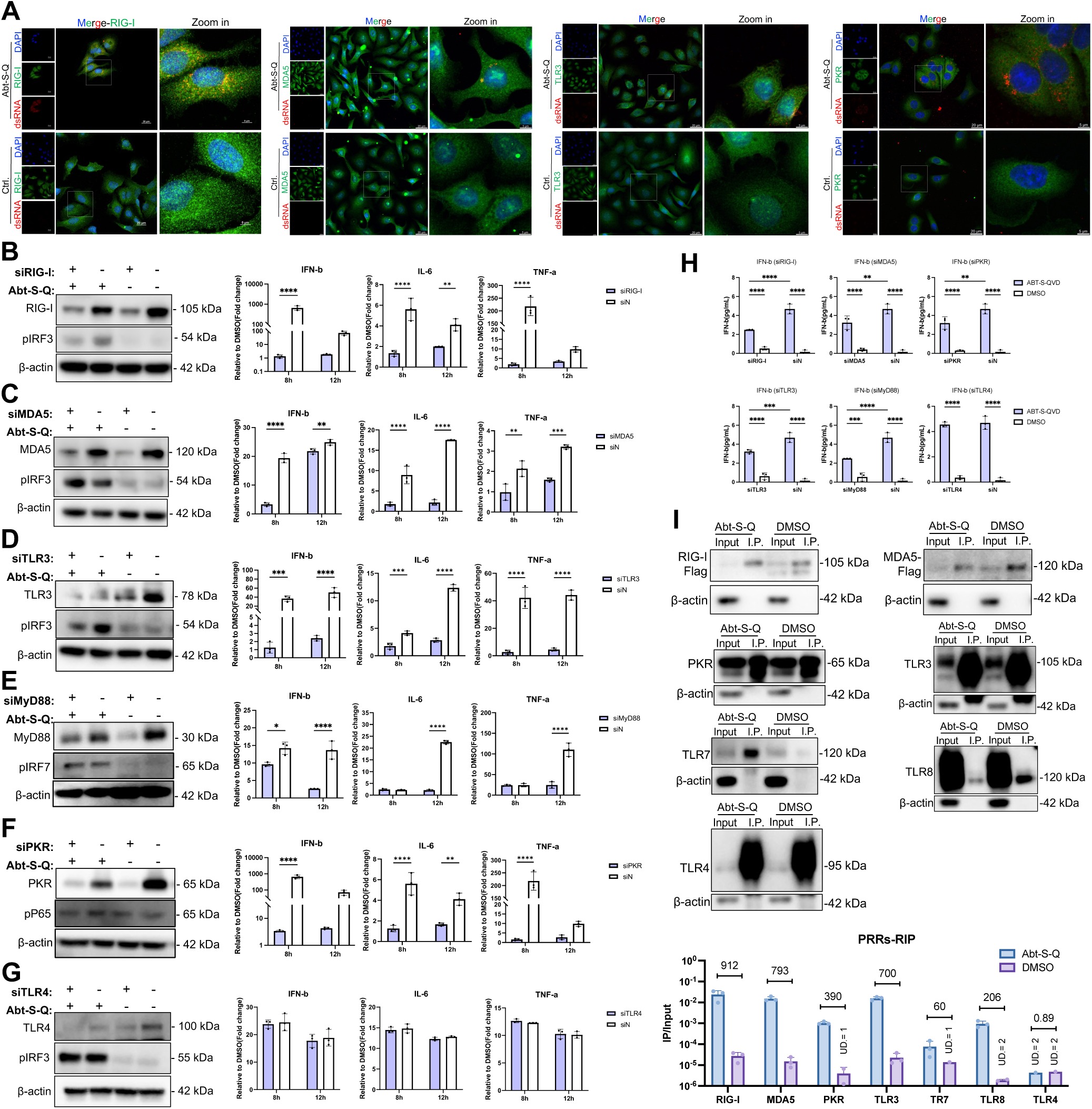
The extruded mtRNA actives innate immunity. (A) A549 cells were treated with Abt-S-Q at 37°C for 90 min followed by IFA for colocalization evaluation of extruded mtRNA and indicated cytosolic PRR molecules. (B-H) A549 cells were transfected with siRNA targeting RIG-I (B), MDA5 (C), TLR3 (D), MyD88 (E), PKR (F) or TLR4 (G) 72 h prior to Abt-S-Q-treatment. Cells were collected for knockdown confirmation by immunoblotting (8 h), and transcription evaluation for IFN-β, IL-6, and TFN-α by RT-qPCR (8 and 12 h). The cell culture medium was collected at 16 h for IFN determination (H). (I) A549 cells were transfected with plasmid expressing Flag-tagged RIG-I, MDA5, PKR, TLR3/7/8/4 24 h prior to Abt-S-Q treatment. The overexpressed PRRs were confirmed in Input and immunoprecipitation (I.P.) fraction. The total mtRNA (Input) and the PRR molecule-bound mtRNA (I.P.) were quantified by RT-qPCR. Data presented as mean ± SD. n = 3 biological replicates. Two-way ANOVA was used. \**p* < 0.05, ** *p* < 0.01, *** *p* < 0.001, **** *p* < 0.0001.

### N6-methyadenosine (m^6^A) on mtRNA is supported by sequencing data and MeRIP assays

In our previous work we discovered that exogenous RNA such as viral RNA could be m^6^A modified by host m^6^A machinery, and thus escape from RIG-I-MAVS-involved innate immunity^23,24,38^. So we wondered if mtRNA also experiences post transcription modification (PTM) of m^6^A in its distinct synthesis site of mitochondrial matrix. To date there’s only one report about METTL17-mediated mtRNA’s m^6^A modification detected by LC-MS/MS^30^, which is not able to locate m^6^A along the RNA strand. Thereby we searched the public database GEO with the keyword “m^6^A” for sequence raw data. There’re 5017 Homo sapiens (human)-related results by the date of Dec.16, 2025, and by adding extra keyword of “MeRIP” or “Nanopore” there’re 783 and 28 accessible datasets, respectively. The raw data of 5 published (GSE29714, GSE55572, GSE231737, GSE185990, GSE132971) and 1 unpublished (GSE296858) m^6^A-related studies, spanning from 2012 to 2025, were downloaded and visualized by software igv^39–43^. In the workspace window m^6^A peaks acquired by MeRIP are presented as two overlapped tracks of input and IP (GSE29714, GSE55572), or one normalized track (GSE231737). Each sample by nanopore has only one track due to the direct sequencing method, but the tracks of YTHDF1/ METTL3-kncokdown sample and the corresponding control were also overlapped to reflect the difference upon m^6^A machinery deficiency. In all 6 datasets m^6^A peaks were demonstrated to scatter along the whole mitochondrial transcript (chrM), including D-loop, mt-mRNA, mt-tRNA and mt-rRNA regions. In three MeRIP datasets based on different cell lines or primary cells, the m^6^A peaks showed relatively conserved location in certain region such as D-loop, ND1-2, COX1-2, ATP8, ATP6, ND4 and CYTB (Figure 4A). The nanopore datasets illustrated m^6^A peaks extensively spreads throughout the polycistronic RNA (Figure 4A). More excitingly we found that METTL3-knockdown consistently reduced m^6^A enrichment in the regions of D-loop, ND1-2, COX2, ND4 and CYTB in both datasets (GSE55572 and GSE185990) by different methods (Figure 4A). Moreover the m^6^A peaks in mRNA of other chromosomes (chr, e.g. chr1, 14 and 20) are presented with much lower data range, like 0-500 for chr1 *vs.* 0-150,000 for chrM in GSE29714, 0-200 for chr1 *vs.* 0-100,000 for chrM in GSE55572, and 0-500 for chr1 *vs.* 0-200,000 for chrM in GSE185990 (Figure 4A and B, *S.*Figure 3A and B), indicating the sufficient sequencing depth and high confidence for mtRNA. In m^6^A antibody-dependent methylated RNA immunoprecipitation (MeRIP), mtRNA of ND2, ND6, CYB, COX1 and ATP8 was detected from the m^6^A antibody-bound IP (Figure 4C), as well as β-actin (ACTB), which is also m^6^A-modified according to the GEO data (*S.*Figure 3C). When a chemical STM2457 was applied to specifically inhibit METTL3, the enrichment of mtRNA dropped significantly, as well as m^6^A-modified nucleus-encoded mRNA such as ACTB, POLRMT and PNPT1, however, the unmodified IFN-β (IFNB1) mRNA is not affected by STM2457 (Figure 4D, *S.*Figure 3C-F). In the context of rVSV-GFP infection, the MeRIP enrichment of ND6 and COX1 was significantly upregulated at 8 HPI, meanwhile the modification of some nuclear-encoded transcripts (ELAC1 and PNPT1) was downregulated. Comparing with the unaffected ACTB, the m^6^A enrichment of IFNA1, IFNB1 and IFITM123 was dramatically enhanced (Figure 4E), suggesting a reprogramming of m^6^A upon innate immunity activation. This findings strongly supports our hypothesis that m^6^A exits on mitochondrial transcripts, among which some are sensitive to the canonical m^6^A machinery, or even the virus-induced m^6^A landscape alternation. In METTL3- and ALKBH5-overexpressing cells mtRNA copies were enhanced and decreased, respectively (Figure 4F), while these canonical m^6^A enzymes didn’t affect the tested mitochondrial enzymes involved in mtDNA-replication and - transcription (TFAM, POLRMT, TEFM), mtRNA-process (TRMT10C, ELAC2), - stability (FASTKD3) and - decay (PNPT1, SUV3L1) (Figure 4G). Thus, it raises the possibility that the canonical m^6^A machinery may functions on the biosynthesis of mtRNA within the mitochondrial matrix, although the underlying mechanism remains to be elucidated.

**Figure 4.**
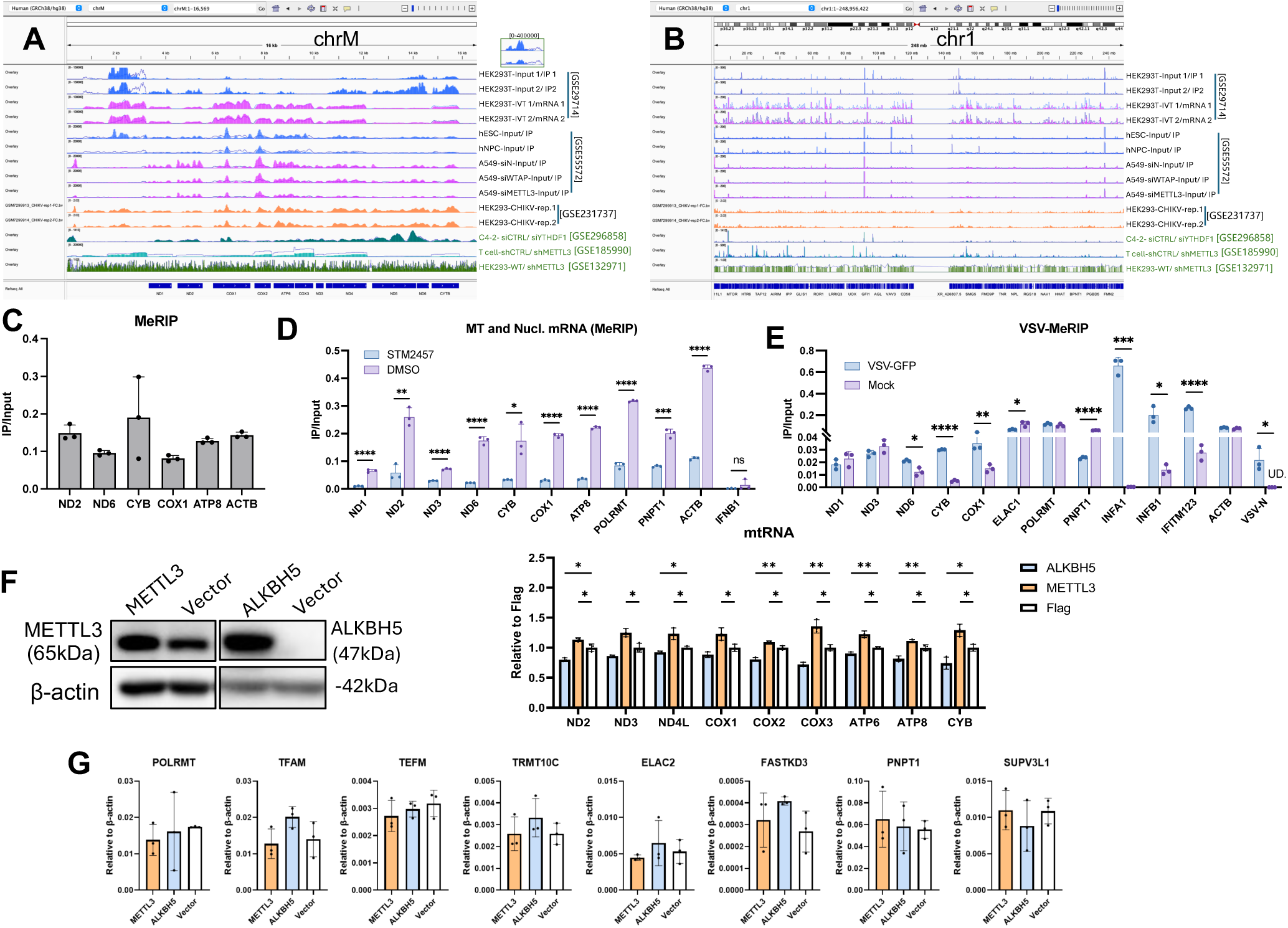
m^6^A modification is detected in mtRNA. (A and B) Human’s m^6^A raw data were searched in Gene Expression Omnibus (GEO) data base (https://www.ncbi.nlm.nih.gov/gds) and downloaded for visualization and alignment across independent studies. The m^6^A peaks on mitochondrial transcripts (chrM) and chrome 1 (chr1) are presented with software igv. The tracks labeled with GEO accession numbers in black were sequenced by MeRIP and the green ones by Nanopore direct RNA sequencing. The tracks of Input or Ctrl. samples are presented with line plot, and the tracks of IP or siRNA-knockdown are presented with bar chart. (C) The mtRNA was enriched with m^6^A-antibody mediated methylated RNA immunoprecipitation (MeRIP) controlled with β-acitn (ACTB). (D) The A549 cells were treated with STM2457 (5 µM) for 2 days prior to the total RNA collection and MeRIP enrichment for m^6^A-modified mtRNA/mRNA quantification. (E) The A549 cells were infected with rVSV-GFP (MOI=0.01), the total RNA was collected at 8 HPI for MeRIP. (F and G) A549 cells were transfected with Flag-tagged METTL3 or ALKBH5 48 h prior to protein and total RNA collection. The overexpression was confirmed by WB, mtRNA (F) and nuclear-encoding mitochondrial enzyme transcripts were quantified by RT-qPCR (G). Data presented as mean ± SD. n = 3 biological replicates. Two-way ANOVA was used. \**p* < 0.05, ** *p* < 0.01, *** *p* < 0.001, **** *p* < 0.0001.

### mtRNA induced innate immunity is modulated by m^6^A machinery

To explore whether the released mtRNA is also substrate of canonical m^6^A machinery in cytoplasm we performed IFA and found colocalization of mtRNA with METTL3, METTL14, FTO, ALKBH5, YTHDF3, or YTHDC1 (Figure 5A). This result implies the possibility that the released mtRNA might be m^6^A-modified in cytoplasm to avoid being recognized by cytoplasmic PRRs, though there’s yet no proof about m^6^A installment on mitochondrial transcripts except for transcription factor B1 (TFB1M)-catalyzed two m^6^2A on 12s rRNA at the position of 936 and 937. We hypothesized the members of canonical m^6^A machinery install or remove mtRNA’s m^6^A either in mitochondrial matrix during the process to facilitate mtRNA maturation, or in cytoplasm upon mtRNA extrusion to modulate innate immune response. When the ALKBH5-, METTL3- and FTO-overexpressing A549 cells were treated with Abt-S-Q, only ALKBH5 enhanced the cytoplasmic mtRNA-induced innate immunity (Figure 5B). The METTL3-overexpression resulted lower cytoplasmic mtRNA upon the Abt-S-Q treatment, while ALKBH5 functioned reversely (Figure 5C). Overexpression of m^6^A reader protein YTHDF3 also prolonged the extruded mtRNA’s retention time in cytoplasm and promoted the innate immune response (Figure 5D). As we noticed that METTL3 overexpression did not suppress mtRNA-induced innate immunity (Figure 5B), which might be complicated by the METTL3-mediated global m^6^A enhancement, resulting promoted translation of specific genes such as IFN-β, TNF-α, and NF-κB^44–48^.

**Figure 5.**
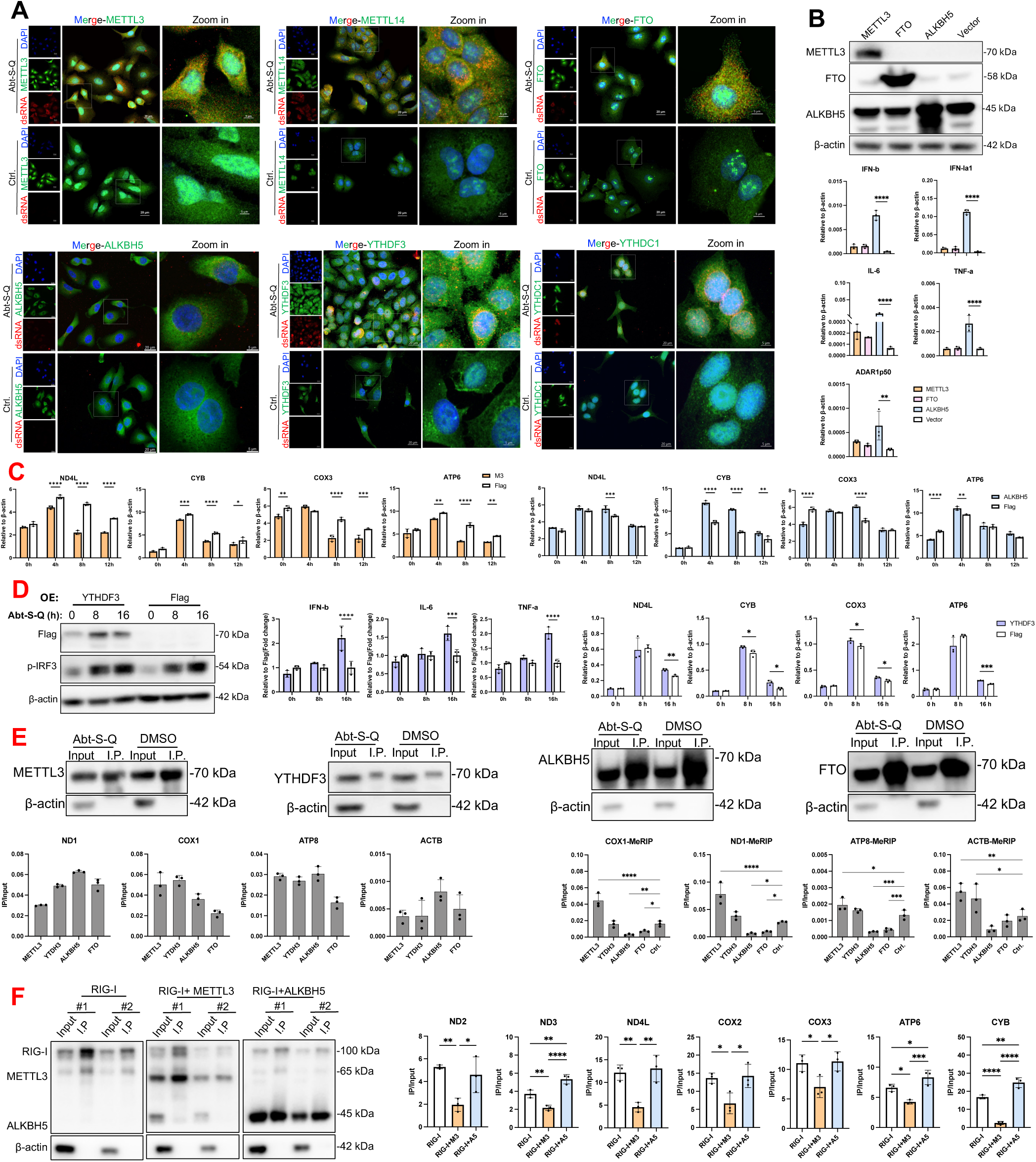
The extruded mtRNA-induced innate immunity is modulated by the classic m^6^A machinery. (A) A549 cells were treated with Abt-S-Q at 37°C for 90 min followed by IFA for colocalization assessment of extruded mtRNA and indicated m^6^A machinery proteins. (B and C) A549 cells were transfected with plasmid expressing METTL3, ALKBH5 or FTO 24 h prior to Abt-S-Q-treatment. The mtRNA-induced innate immunity was evaluated with transcription of IFN-β, IL-6, TFN-α, and IFN-induced ADAR1p150 at 8 h post treatment (B), and the extruded mtRNA copies were determined from mitochondria-excluded cytoplasm at 0, 4, 8, and 12 h by RT-qPCR (C). (D) HEK293T cells were transfected with plasmid expressing YTHDF3 or vehicle control 24 h prior to Abt-S-Q-treatment. The innate immunity was evaluated with transcription of IFN-β, IL-6, and TFN-α, and the extruded mtRNA copies were determined from mitochondria-excluded cytoplasm at 0, 8, and 16 h. (E) A549 cells overexpressing the indicated m^6^A machinery proteins were treated with Abt-S-Q for 2 h and then were subjected to UV-cross link, followed by RIP (bottom left panels) and MeRIP (bottom right panels) assays. The protein- or m^6^A antibody-bound mtRNA was quantified by RT-qPCR. (F) A549 cells were co-transfected with plasmids expressing Flag-tagged RIG-I and 6×His-tagged METTL3/ ALKBH5, controlled with single RIG-I-transfection, 24 h prior to Abt-S-Q-treatment. The overexpressed RIG-I and METTL3/ ALKBH5 were determined in Input and I.P. samples by immunoblotting (left panel), the RIG-I-bound mtRNA was pulled down by Flag antibody-conjugated magnetic beads and quantified by RT-qPCR (right panel). Data presented as mean ± SD. n = 3 biological replicates. Two-way ANOVA was used. \**p* < 0.05, ** *p* < 0.01, *** *p* < 0.001, **** *p* < 0.0001.

Furthermore, the introduced expression plasmids can activate innate immunity through cGAS-STING signaling pathway^49^. To exclude these interfering factors, the protein-RNA binding assay (RIP) and MeRIP assay were carried out in the METTL3-, YTHDF3-, ALKBH5- or FTO-overexpressing A549 cells after Abt-S-Q treatment. All four m^6^A machinery proteins are capable of binding the cytosolic mtRNA with varying affinities (IP/Input ratios). Furthermore, following 2 hours of cytosolic co-exposure, m^6^A levels on mtRNA were altered. Specifically, METTL3 significantly enhanced the m^6^A enrichment, consistent with its effect on nuclear-encoded mRNA (ACTB), whereas FTO and ALKBH5 reduced m^6^A levels (Figure 5E). These findings suggest the possibility that the extruded mtRNA undergoes “secondary modification” in the cytoplasm by m^6^A machinery, analogous to its activity on viral RNA. Co-overexpression with METTL3 significantly reduced RIG-I-bound mtRNA compared with the single RIG-I-overexpression, and as expected the RIG-I-bound mtRNA was enriched when ALKBH5 was co-overexpressed (Figure 5F). However not all detected mtRNA copies exhibited increased affinity for RIG-I in ALKBH5-overexpressing cells, suggesting the disequilibrium of m^6^A location across mitochondrial transcripts (Figure 4A), analogous to the location-dependent effects of m^6^A on nuclear-encoded transcripts^50,51^. Thus our findings enlighten the possibility that extruded mtRNA is potential substrate of canonical m^6^A machinery when they encounter in the cytoplasm, and the dynamic modification or m^6^A-based binding by reader proteins masks this “exogenous” dsRNA from being recognized by RIG-I, or modulates its degradation, for the benefit of homeostasis in cytoplasm.

## Discussion

Multiple studies reported that mtDNA is released through either oligomerized Bax macropore or VDAC1 oligomers-associated mPTP depending on the stress^2,52–54^, however, mtRNA, particularly mt-dsRNAs, was reported to through Bax/Bak macropores and mitochondrial outer membrane permeabilization (MOMP)^55^. Bcl-2 and Mcl-1 cooperatively inhibit the Bax/Bak channel, thereby preventing MOMP. Here, we used chemical inhibitors to establish an aberrant Bax/Bak activation model. Abt-737 and S63845 specifically inhibit Bcl-2 and Mcl-1, respectively, and Q-VD-Oph is a pan-caspase inhibitor that delays caspase pathway activation and apoptosis induced by Bax/Bak channel opening, thereby extending the observation window^3^. The combined treatment with these inhibitors resulted in significantly increased mtRNA copies in mitochondria-depleted cytosol portion, indicating enhanced opening of Bax/Bak pores and mtRNA release into the cytosol, where multiple PRRs function to recognize these “exogenous” RNA and DNA molecules to initiate innate immune responses, which is characterized by elevated transcription of type I IFNs and inflammatory factors, as well as the transcripts of ADAR1p150, a product of alternative splicing induced by type I IFNs and a proof for the IFN secretion^56,57^.

Dengue virus and HSV introduce mtDNA extrusion restrain the viral propagation by activating cGAS-STING or RNA Polymerase III/RIG-I Pathway^14,32^. To our knowledge mtRNA hasn’t been mentioned upon viral infection by the time we write this manuscript. In VSV-infection we observed the host’s mitochondrial network disruption, similar to mitochondrial dysfunction observed during HSV infection and mtDNA extrusion. Moreover, the increased mtRNA was detected over time by RT-qPCR from the infected A549 cytoplasm. VSV and other NNS virus infection activate RIG-I-MAVS axis to launch type I IFN response^23^. Moreover, the interactions between MAVS and other mitochondrial membrane proteins, such as Dynamin-related protein 1 (DRP1), Prohibitins (PHBs complexes on mitochondrial inner membrane), VDAC1 and HK2, also contribute to MMP maintenance, dynamics and permeability of the mitochondria^58–61^. In this study we unexpectedly discovered that VSV resulted in obvious MAVS degradation and HK2 loss in both membrane- and cytosol-associated portions, accompanied by mtROS accumulation and MMP drop at the middle-to-late stage of infection (12-20 HPI). Therefore, we hypothesize that VSV-induced innate immune response may triggers MAVS degradation, possibly via the K-48 linked ubiquitination–proteasome pathway^62,63^, thereby disrupting the functional interplay of MAVS-HK2-VDAC1 on the OMM, and facilitated aberrant activation and oligomerization of VDAC1. Furthermore, accumulated mtROS may promote the redox-sensitive protein VDAC1 to undergo oligomerization^64^, and ultimately leads to mPTP activation. In lupus-like disease such an opening of mPTP and transportation of mtNAs across the inner and outer mitochondrial membranes under oxidative stress conditions were also observed^2^. Besides VDAC1, Bax was proved to participate VSV-resulted mtNAs extrusion. Bax was reported to translocate to OMM for Bax/Bak macropore opening during IAV and Sendai virus infection, and MAVS-mediated IRF3 phosphorylation is also essential for Bax/Bak channel opening and the subsequent mtDNA release^13,65^. Thus the virus activated RIG-I-MAVS signaling pathway promoted the mtRNA extrusion through multiple mechanisms, which need to be comprehensively illustrated in the future. Conversely, when mtRNA extrusion is pre-established due to other factors, the subsequent viral infection is markedly suppressed.

Early studies reported that released mtRNA is recognized by cytosolic PKR^17,18^ and MDA5^15,20^, as well as endosomal TLR3^20^, leading to innate immune activation. Most recently Doke et al. reported that mtRNA can activate the RIG-I-MAVS signaling pathway^21^. Unlike nuc-mRNA and many viral mRNA, which are protected by N7-methylguanosine (m^7^G) cap structure, mtRNAs possess only a 5′-triphosphate (5′-ppp) group without m^7^G cap, just like some virus genomic and antigenomic RNA^23,55^. Based on our early observation on the viral 5′-ppp RNA-activated RIG-I-MAVS axis, we hypothesize that once released into the cytoplasm the mtRNA is highly likely to be recognized by RIG-I through high affinity binding between uncapped 5′-ppp and C-terminal domain (CTD) of RIG-I^23,24,66^. In the Abt-S-Q-induced model, the released mtRNA was bound by not only RIG-I and MDA5, but also TLR3/7/8 and PKR molecules. It has been reported that mitochondrial 16S rRNA-derived fragments can activate TLR8 when introduced by transfection^67^, and in certain circumstance (e.g. the *Irgm1* gene deletion) mtRNA can be delivered to endososomal compartment via canonical mitophagy related mechanisms, and activates TLR7^68^. TLR7/8 are though not the predominant RNA-sensing PRRs in A549, however, the overexpressed molecules can bind mtRNA with lower efficiency compared to RLRs and TLR3. But how cytoplastic mtRNA is delivered endosomal TLRs remains unclear and warrants further investigation; potential mechanisms may include exosome-mediated trafficking or endocytosis/vesicular transport.

Our previous work revealed that the affinity of RIG-I for viral RNA is highly influenced by m^6^A modifications on the ligand RNA^23,24,38^. The proof of m^6^A modified mammalian mtRNA is still lacking and ambiguous, which might be due to the technical challenge or focus bias in research, except for the most recent evidence by LC-MS/MS^30^. However, research groups have detected m^6^A peaks in Arabidopsis thaliana’s and cauliflower’s mtRNAs^69,70^. In Arabidopsis, over 350 m^6^A sites (∼4.6-4.9 sites per transcript) have been mapped in mtRNA, with an overall m^6^A level substantially higher than that observed in nuclear transcripts, and the m^6^A motif sequences in plant mtRNA are similar to those in nuc-mRAN^70^. Because of the shortage of direct m^6^A-sequencing result we applied the raw data of m^6^A MeRIP and Nanopore direct RNA sequencing from GEO database (https://www.ncbi.nlm.nih.gov/geo/). The sequencing datasets exhibited even higher coverage and sequencing depth of m^6^A in mtRNA than in nuc-mRNA, consistent with the observations in Arabidopsis^70^. The following MeRIP assays further support the presence of m^6^A in mtRNA, and indicate the modification level can be modulated response to the METTL3 inactivation and VSV-infection, similar to nuclear-encoded mRNA.

The m^6^A methyltransferases and demethylases predominantly locate in nucleus and can be induced into cytoplasm by various stresses such as RNA virus infection, to modify or bind viral RNA for innate immunity escape and more efficient replication of the invading virus^24,71,72^. In this study we also revealed that the extruded mtRNA colocalized with m^6^A machinery including enzymes and binding proteins. The overexpression of either METTL3 or ALKBH5 significantly affected the cytosolic retention time of mtRNA. The enhanced YTHDF3 also prolonged the life time of mtRNA and thus made the IFN response sever. Not all mitochondrial transcripts (7/14) showed altered binding to RIG-I upon METTL3 perturbation, and even fewer exhibited modest and less pronounced effects upon ALKBH5 overexpression (3/14). We therefore speculate that m^6^A-mediated RIG-I recognition depends not only on the abundance but also on the positioning of m^6^A marks, for example through m^6^A-dependent secondary structure remodeling (e.g., dsRNA formation) and reader protein recruitment, as reported for cellular and certain viral RNAs, thereby modulating RIG-I affinity^73–75^.

In summary, we for the first time demonstrated that aberrant Bax/Bak channel open-induced mtDNA release is accompanied by mtRNA through experimental methods, and illustrated that VSV infection is capable of resulting in mtRNA extrusion, probably mediated by MAVS degradation and mitochondrial membrane integrity disruption, whose detailed mechanism needs to be clarified by further and deeper dig. These two *in vitro* models of mtRNA extrusion, as well as the one based on SUV3/PNPT1 deficiency, will be useful to explore therapeutic methods for the benefit of autoimmune and degenerative diseases, in which mtRNA accumulates in cytoplasm abnormally due to genetic or acquired etiology. Our findings also raise the possibility that extruded mtRNA serves as a substrate for cytoplasmic m^6^A machinery, thereby attenuating its recognition by RIG-I and preventing excessive innate immunity.

## Limitations of the study

This study had several limitations. Firstly, the mechanism of VSV-induced mtRNA extrusion is based on the WB and mitochondrial functional assay, however, the speculated MAVS-HK2-VDAC1 pathway needs to be illustrated more directly and explicitly. Secondly, although multiple independent m^6^A datasets in GEO, along with our own MeRIP analyses, consistently support the existence of m^6^A modifications on mtRNAs, we lack direct mtRNA m^6^A-sequencing data generated in our own experimental system, and it remains unclear whether canonical m^6^A machinery functions in the mitochondrial matrix as it does in the nucleolus and cytoplasm.

Thirdly, extruded mtRNA, together with mtDNA, has been shown to contribute to the VSV-induced innate immune response, however, whether mtRNA-RIG-I interaction represent the predominant regulatory axis during viral infection remains uncertain due to the involvement of multiple PRRs. This will be addressed in our future studies.

## RESOURCE AVAILABILITY

### Lead contact

Further information and requests for the resources and reagents should be directed to and will be fulfilled by the lead contact, Mijia Lu

### Materials availability

- All unique/stable reagents generated in this study are available from the lead contact without restriction.
- The siRNA oligonucleotide sequences were provided in Table S1, and primer sequences of target genes were provided in Table S2.

### Data and code availability

- This paper does not report the original code.
- Any additional information required to reanalyze the data reported in this work paper is available from the lead contact upon request.

## ACKNOWLEDGMENTS

This work is supported by grants from the National Natural Science Foundation of China (No. 32270148), the Joint Funds of the National Natural Science Foundation of China (No. U22A20342), and the Major Program of the National Natural Science Foundation of China (No. 82595941). The authors thank all the members of our laboratory for helpful discussion, and acknowledge the support from the Scientific Research Center, Hangzhou Medical College.

## AUTHOR CONTRIBUTIONS

Q.H.: investigation, data collection, formal analysis and validation. Y.-f.Z., S.-x.W., X.-p.Z., F.-y.W.: data collection and analysis. T.F.: Project administration. Y.-q.C.: writing-original draft preparation., J.-x.L.: funding acquisition, supervision. M.-j.L: conceptualization and methodology, writing-reviewing and editing, funding acquisition, supervision. All authors contributed to the article and approved the final manuscript.

## SUPPLEMENTAL INFORMATION

Document S1. Figures S1–S3, Tables S1 and S2.

## STAR★METHODS

### KEY RESOURCES TABLE

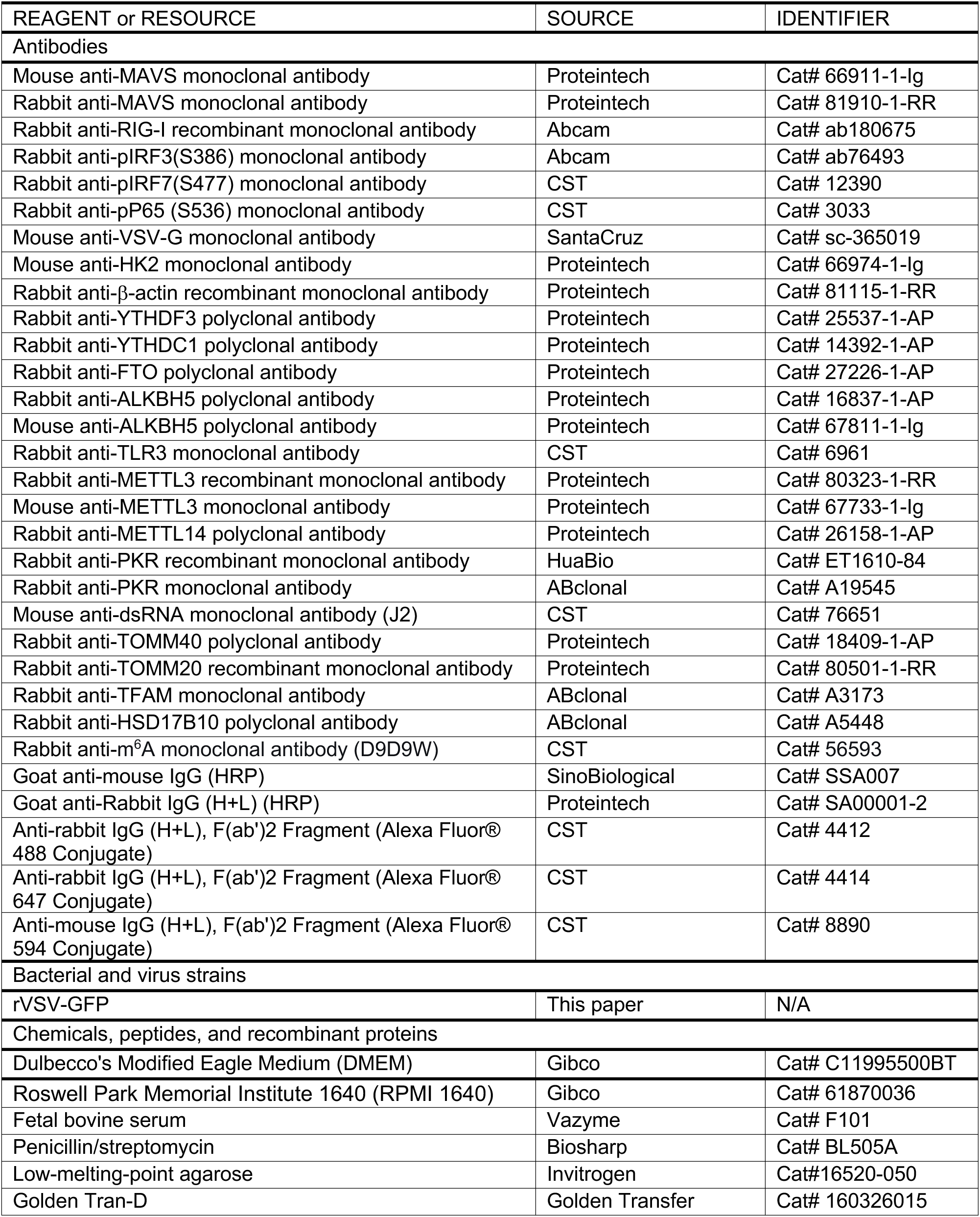
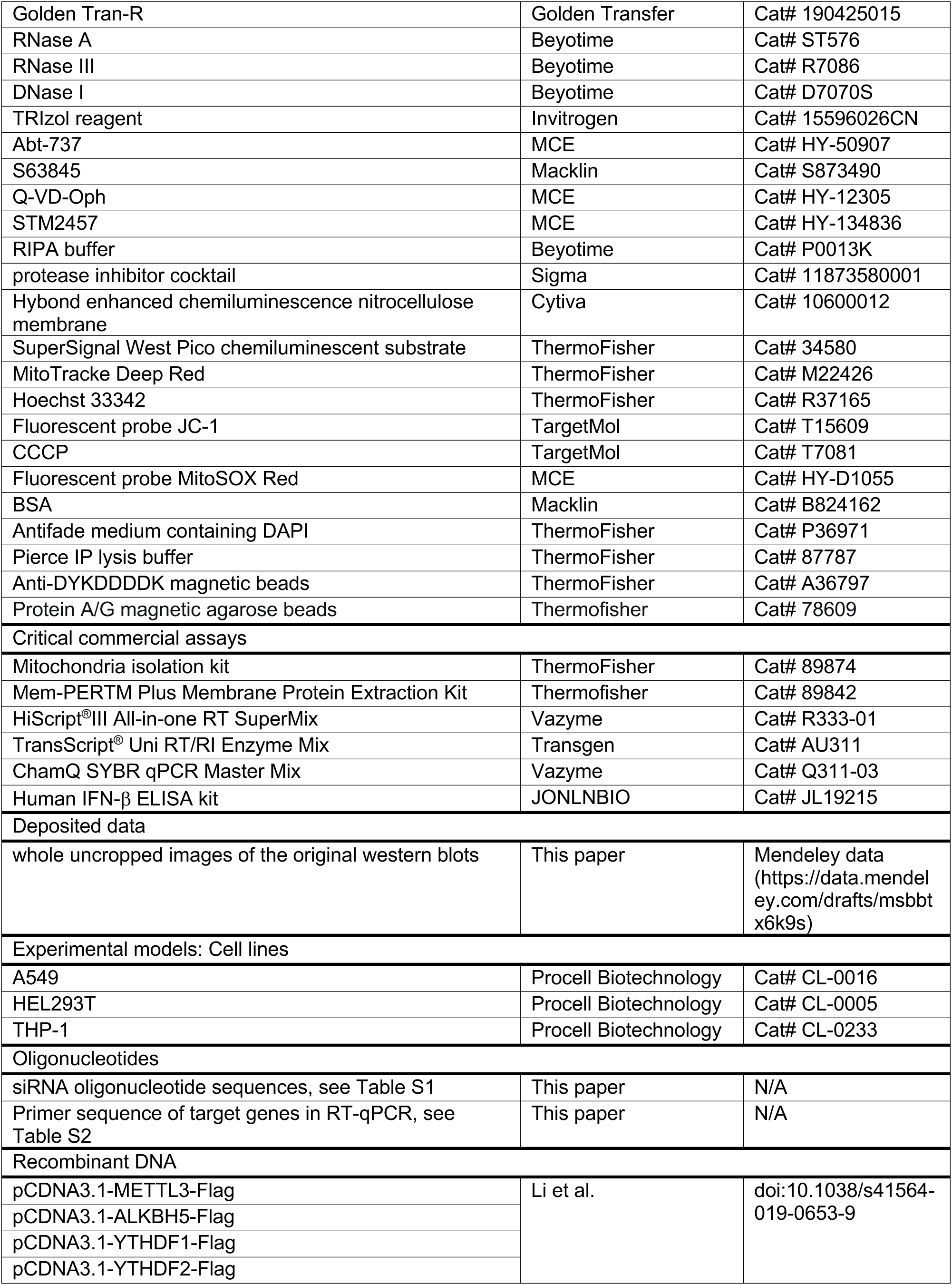
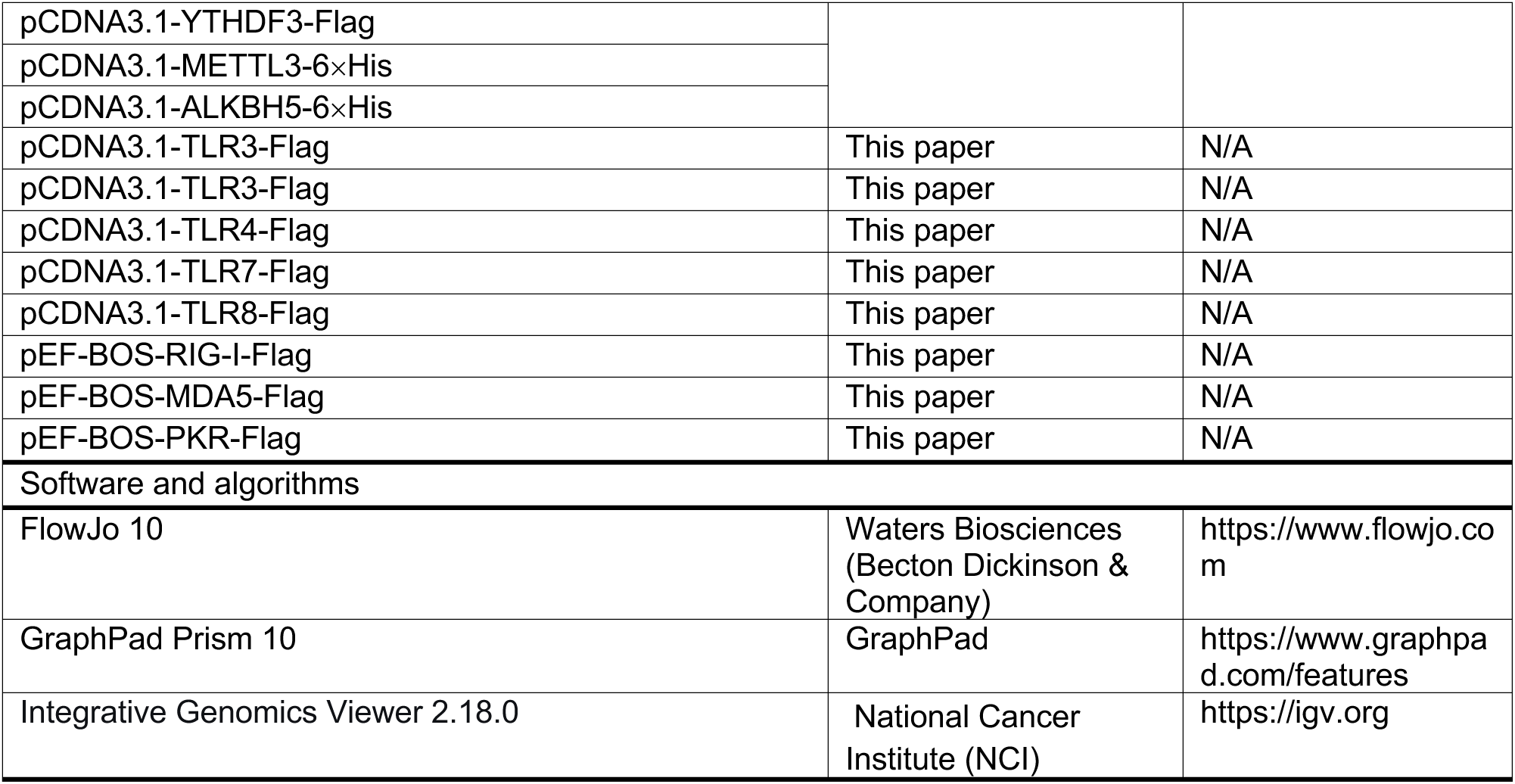

### METHOD DETAILS

#### Biosafety statement

All experiments with infectious VSV were conducted under biosafety level 2 (BSL2) at Hangzhou Medical College using standard operating procedures and were approved by the BSL2 Advisory Group and the Institutional Biosafety Committee (IBC).

#### Cell line

A549 and HEK293T cell lines were purchased from Procell Biotechnology Co., Ltd. Wuhan, China, and were cultured in Dulbecco’s Modified Eagle’s medium (DMEM, Gibco) supplemented with 10% fetal bovine serum (FBS, Sigma) and 1% penicillin/streptomycin (Biosharp). The human monocytic cell line THP-1 was cultured in Roswell Park Memorial Institute (RPMI) 1640 medium (Gibco) supplemented with 10% FBS and 1% penicillin/streptomycin. All the cells were cultured in 37°C incubator with 5% CO_2_.

#### Chemical drugs-treatment in vitro

To induce mitochondrial nucleic acids (mtNA) extrusion from mitochondria into cytoplasm, A549 cells were treated with a mixture of chemical drugs containing Abt-737 (5 μM, MedChemExpress, MCE), S63845 (25 μM, Macklin) and Q-VD-Oph (20 μM, MCE) in a 37°C incubator for the indicated time, and the combination of these three drugs will be referred as Abt-S-Q hereinafter.

#### Virus and infection

The recombinant Vesicular Stomatitis Virus expressing GFP (rVSV-GFP) stock used in this study was rescued with plasmid containing full length genomic cDNA of VSV Indiana strain, and assistant plasmids expressing VSV N, P and L proteins, through reverse genetic operation.

Cells were seeded in culture flask or multi-well plate and incubated at 37°C with 5% CO_2_ 16∼24 h prior to the infection. The cell’s culture medium was then removed and the VSV dilution was added to inoculate the cells at a multiplicity of infection (MOI) of 0.005 or 0.01 as indicated with constant and gentle shaking for 1 h. The inoculum was then replaced with culture medium containing 2% FBS and the cells were incubated at 37°C until the indicated harvest time points.

#### Virus titration by plaque assay

Confluent Vero cells in 6-well plates were infected with 10-fold serial dilutions of rVSV-GFP in FBS-free DMEM. After absorption for 1 h at 37°C, the virus inoculum was replaced with the 2 mL overlay containing low-melting-point agarose (0.25% w/v), 2% FBS, 0.12% sodium bicarbonate, 100 mg/mL streptomycin, 100 U/mL penicillin, 25 mM HEPES (pH 7.7), and 2 mM L-glutamine. After incubation at 37°C for 40 h, cells were fixed with 4% neutral buffered formaldehyde for 2 h. The overlays were removed, and the plaques were visualized by crystal violet staining. The virus-formed plaques in one well were calculated as plaque-forming unit per mL (PFU/mL).

#### mtRNA purification

The HEK293T cells harvested from 4× 15 cm culture dishes were used to isolate mitochondria as described above. The resulted mitochondria pellet was resuspended in 50 μL reaction system containing 10 mM Tris-HCl (pH 7.4), 15 mM NaCl, 1 mM EDTA, 10 μg RNase A (Beyotime) and 5 U RNase III (Beyotime), incubated at 37°C for 20 min and then lysed with TRIzol reagent (Invitrogen) for mtRNA purification. To remove mtDNA, the mtRNA was treated with 20 U of DNase I (Beyotime) at 37°C for 20 min and purified with TRIzol again to remove DNase. The final mtRNA was quantified for mass concentration on a Nanodrop, and evaluated of mtDNA-contamination by RT-qPCR and direct qPCR.

#### Plasmids- and RNA-transfection

Plasmids expressing Flag-tagged METTL3, ALKBH5, YTHDF1, 2, 3, RIG-I, MDA5, or empty vector was transfected into cells using reagent Golden Tran-D (Golden Transfer Science and Technology Co.Ltd. Changchun China) according to the manufacturer’s instructions. Briefly, ninety percent confluent A549 or HEK293T cells in a 24-well plate were transfected with 0.5 μg of plasmid 24 h prior to the following experiment. To knockdown genes siRNAs against RIG-I, MDA5, TLR3, MyD88, PKR or non-targeting negative control siRNA (custom synthesized by Tsingke Biotech Co., Ltd. Beijing, China, sequences were listed in Table S1) were used to transfect cells in 24-well plate (15 pmol/well) by Golden Tran-R (Golden Transfer) following the manufacturer’s instructions 72 h prior to the following experiment. For mtRNA, Lipofectamine 3000 (Thermofisher) was used as transfection reagent to introduce mtRNA into A549 cells with the indicated amount per well.

#### Isolation of cytoplasmic and mitochondrial portions from cultured cells

Cells were collected by gentle scraping and transferred into tubes for the subsequent mitochondrial removing using Mitochondria isolation kit (ThermoFisher) according to the manufacturer’s instructions. Briefly, cell pellet was acquired by centrifugation at 850 ×g for 2 min at 4°C, treated with reagent A on ice for exactly 2 min, reagent B on ice for 5 min, and then reagent C before the centrifugation at 850 ×g for 2 min at 4°C to separate cytosol from mitochondrial fractions. The final cytosol portion was subjected to protein sample preparation or RNA extraction.

#### Isolation of cytosolic and membrane-associated protein

The infected cells in 6-well plate (10^6^ cell/ well) were collected by gentle scraping and cell pellet was harvested by centrifugation at 300 ×g for 5 min. The cell pellet was used to isolate cytoplasmic and membrane-associated proteins with Mem-PERTM Plus Membrane Protein Extraction Kit (Thermofisher) according to the manufacturer’s instructions. Briefly, cell pellet was washed with cell wash solution twice, permeabilized at 4°C for 10 min, and centrifuged for 15 min at 16,000 ×g to remove cytosolic proteins in the supernatant. The pellet was resuspended, lysed at 4°C for an extra 30 min and centrifuged at 16,000 ×g for 15 min at 4°C to collect the membrane-associated proteins in the supernatant.

#### Reverse transcription-Real time PCR (RT-qPCR)

To quantify the transcription of nuclear genes, the total RNA was extracted from cell lysate, or mitochondria-depleted cytosol portion with TRIzol reagent at indicated time points. For nuclear gene’s transcripts (mRNA) determination the cDNA pool was generated with random primer (HiScript^®^III All-in-one RT SuperMix, Vazyme) as templates. For mitochondrial gene’s transcripts (mt-mRNA) quantification, the cDNA pool was generated with anchored oligo (T)_23_ (custom synthesized by Tsingke Biotech) and TransScript^®^ Uni RT/RI Enzyme Mix (Transgene). The qPCR was performed in ChamQ SYBR qPCR Master Mix (Vazyme) using the indicated primer pairs with an internal control of *β-actin*. All the primer pairs used in RT-qPCR were listed in Table S2.

#### Immunoblot assay (Western blot, WB)

At each indicated time point cells were lysed in RIPA buffer (Beyotime Biotechnology) supplemented with protease inhibitor cocktail (Sigma-Aldrich) on ice and denatured in 5× loading buffer at 98°C. Proteins were separated by 12% SDS-PAGE and transferred to a Hybond enhanced chemiluminescence nitrocellulose membrane (Cytiva) in a Mini Trans-Blot electrophoretic transfer cell (Bio-Rad). The blot was probed with the indicated primary antibody overnight at 4°C, followed by horseradish peroxidase (HRP)-labeled secondary antibody incubation for 1 h at room temperature (R.T.), and development with SuperSignal West Pico chemiluminescent substrate (ThermoFisher).

#### Enzyme-linked immunosorbent assay (ELISA)

At 16 h post Abt-S-Q treatment, the culture medium of the treated A549 was collected and clarified by centrifugation at 12,000 rpm for 5 min at 4°C. The supernatant was then subjected to IFN-β quantification by commercial ELISA kit (JONLNBIO) following the manufacturer’s instructions. In brief, 100 μL clarified culture medium or standard reagent were added into capture antibody-coated microwells, and incubated at 37 °C for 1 h. Then the biotin-conjugated detecting antibody, Streptavidin-HRP reagent, and TMB substrate were added in order. After the stop solution was added the microplate was read for optical density (OD) values at 450 nm. The concentration of IFN-β (pg/mL) was calculated according to the standard curve.

#### Mitochondrial dynamics imaging

For mitochondrial dynamics imaging the culture medium of infected A549 cells was replaced with fresh culture medium containing 100nM MitoTracker Deep Red (ThermoFisher, M22426) and Hoechst 33342 (ThermoFisher) at 8 HPI, and cells were incubated in 37°C incubator for another 60 min. The staining medium was then replaced with culture medium and cells were subjected to imaging using confocal microscope (Leica, STELLARIS).

#### Mitochondrial membrane potential (MMP, Δψm) determination

The fluorescent probe JC-1 (TargetMol) enters mitochondrial matrix and forms polymer that emits strong red fluorescence at the status of a relatively high MMP for normal mitochondria, but can’t aggregate into a polymer, thus producing green fluorescence, as MMP decreases during mitochondrial stress. The reagent was used to determine MMP of cells according to the product instruction. Briefly, VSV (MOI=0.01)-infected cells were harvested at 12 and 20 HPI, and CCCP (TargetMol) was used as a positive control by incubating with cells at 10 μM for 30 min. Cells were harvested by trypsin-digestion for single cell suspension and washed with phosphate buffered saline (PBS) for three times. The cell resuspension was incubated with 1 μM JC-1 at R.T. for 30 min avoiding light, washed for 3 times and resuspended with PBS for flowcytometry. The data were analyzed with software FlowJo.

#### Mitochondrial reactive oxygen species (mtROS) quantification

The fluorescent probe MitoSOX Red (MCE), which specifically targets mitochondria and enters the organelle, was used to determine mtROS according to the product instruction. Briefly, VSV-infected (MOI=0.01) or mock-infected A549 cells were harvested at 8 and 12 HPI by trypsin-digestion for single cell suspension and washed with PBS for once. The cell resuspension was stained with 1 μM MitoSOX Red in PBS at 37°C for 30 min avoiding from light, and then washed with PBS for twice to remove unbound MitoSOX Red. The PBS-resuspended cells were subjected to flow cytometry and the data were analyzed with software FlowJo.

#### Immunofluorescence assay (IFA)

A549 cells were seeded in 8-well chamber slide and allowed to attach overnight. Having been treated with chemicals or transfected with expression plasmid, cell culture medium was removed, cells were washed with PBS and fixed with paraformaldehyde (4% v/v in PBS) for 30 min at R.T. The fixed cells were then successively permeabilized with 0.5% (v/v) Triton-X100, blocked with bovine serum albumin (BSA, 1%, v/v, Macklin, B824162), incubated with indicated primary antibody overnight at 4°C and fluorescent dye-conjugated secondary antibody (Table 3) at R.T. for 1 h, and mounted with antifade medium containing DAPI (ThermoFisher). The slide was observed and imaged under a confocal microscope (Zeiss, LSM710; Nikon, AX R with NPSARC; Leica, STELLARIS).

#### RNA immunoprecipitation (RIP) assay

A549 cells in 6 cm culture dish were transfected with pcDNA3.1-Flag or - 6×His vectors expressing RIG-I, MDA5, TLR3/7/8 or m^6^A machinery proteins 24 h prior to Abt-S-Q treatment. Four or 8 hours after the treatment, cells were washed with pre-cold PBS, UV-cross linked at the dose of 200 J/cm^2^ (in the case of RIG-I, METTL3, YTHDF3, ALKBH5 and FTO-targeting RIP) and scraped off the culture dish for cell pellet collection by centrifugation at 4,000 rpm for 5 min at 4°C. The cell pellet was lysed by Pierce IP lysis buffer (ThermoFisher) on ice and clarified by centrifugation at 12,000 rpm for 10 min at 4°C. Two tenth of the supernatant was conserved as Input for WB sampling and RNA quantification, respectively, and the rest was subjected to IP incubation with anti-DYKDDDDK magnetic beads (ThermoFisher) overnight at 4°C, followed by PBS wash twice. One tenth of the resulting beads was conserved for WB sampling as I.P. and the rest was used for RNA extraction followed by mtRNA quantification by RT-qPCR.

#### m^6^A GEO data analysis

There’re massive curated gene expression DataSets, including m^6^A raw data acquired by various sequencing technologies, stored in Gene Expression Omnibus (GEO) database (https://www.ncbi.nlm.nih.gov/gds). We searched with keywords “m^6^A AND Homo sapiens” in GEO DataSets and refined the search results with extra keyword like “MeRIP” or “Nanopore” to collect data by methylated RNA immunoprecipitation, or nanopore direct-RNA sequencing, respectively. The .BED for MeRIP-derived data and .BW files for Nanopore-derived data focusing on regulatory functions of m^6^A machinery, were downloaded as the original data source. The m^6^A landscape was visualized by opening .BED and .BW files in software Integrative Genomics Viewer (igv, v2.18.0), with the reference of human genome (GRCh38/ hg38). In each independent study the tracks of each Input and the corresponding IP were overlapped and presented in the same data range scale (peaks’ sequencing depth).

#### m^6^A methylated RNA immunoprecipitation (MeRIP)

The total RNA was purified from cell lysate using TRIzol reagent. To modulate m^6^A modification, the METTL3 inhibitor STM2457 (MCE) was applied for pretreatment at the dose of 10 μM, or the indicated expression vector was transfected into cells, 24 h prior to the RNA purification. To allow the exposure of mtRNA to the cytoplasmic proteins, the Abt-S-Q was applied 2 h prior to the RNA collection. One microgram of the m^6^A monoclonal antibody (CST) was conjugated on 15 μL protein A/G magnetic agarose beads (Thermofisher) by 1 h incubation at room temperature. The m^6^A antibody-bound beads were then washed with PBS (containing 150 mM NaCl), blocked with 1% BSA, and then incubated with 40 μg total RNA in the presence of RNase inhibitor at 4°C overnight. The RNA-bound beads were washed and used for RNA purification with TRIol reagent. The resulting RNA (IP) and Input RNA were subjected to RT-qPCR for mtRNA and nuclear-encoded transcripts quantification. The ratio of IP and Input was calculated as enrichment m^6^A-modified RNA.

### QUANTIFICATION AND STATISTICAL ANALYSIS

Two-tailed Student’s t test, One-way ANOVA and Two-way ANOVA analysis were performed to assess the statistical significance of differences between groups, \**p* < 0.05, ** *p* < 0.01, *** *p* < 0.001, **** *p* < 0.0001. Data are presented as the mean ± standard deviation (SD).

