## Supplementary Figures for "The mitochondrial RNA extrusion-induced innate immunity is regulated by N6-methyladenosine machinery"

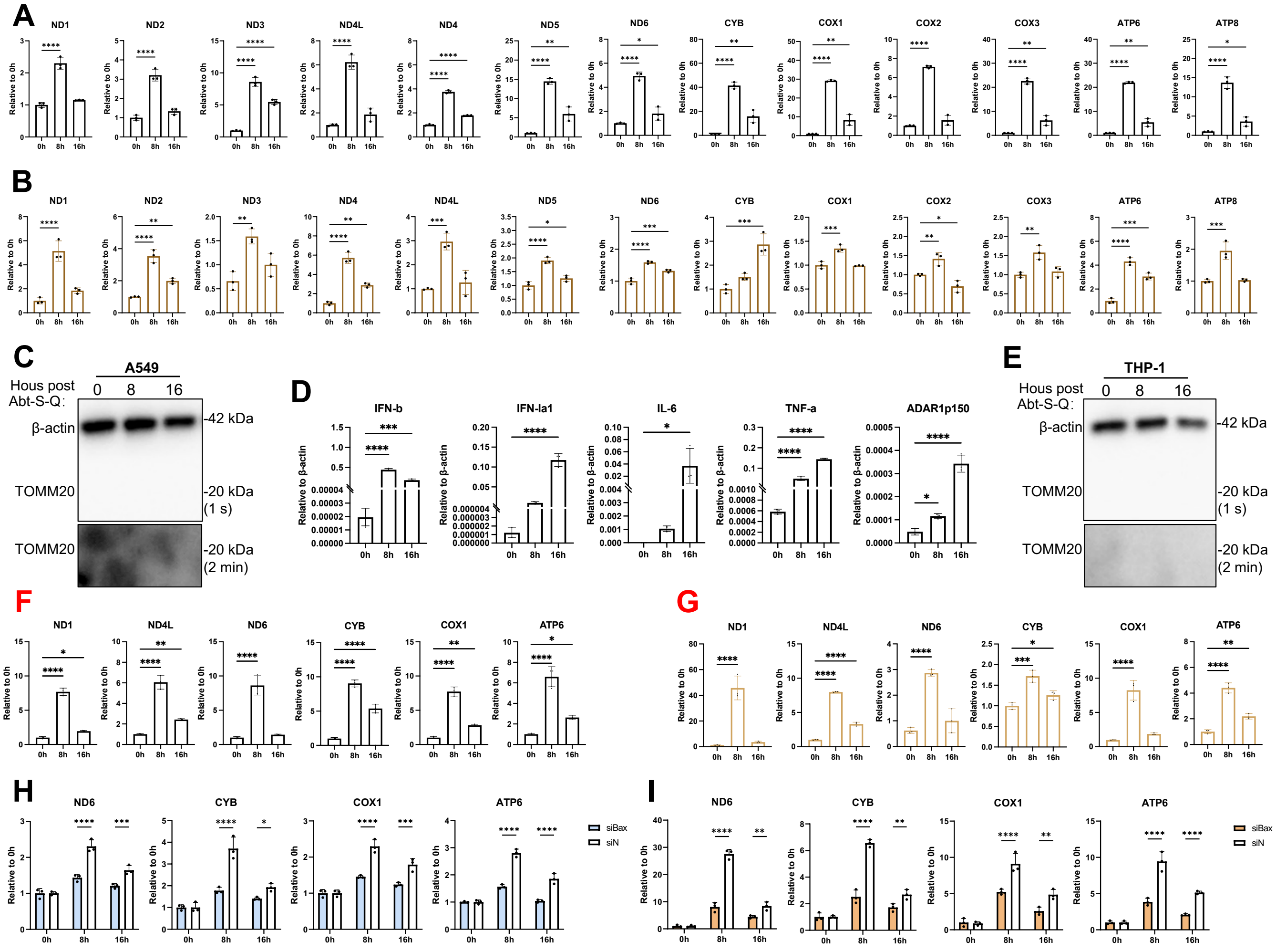


***Supplementary*** **Figure 1. The chemical treatment induces mtRNA extrusion.**

(A-C) A549 cells were treated with Abt-S-Q for 0, 8 and 16 h, mtRNA (A) and mtDNA (B) copies were quantified with RT-qPCR and qPCR, respectively, the mitochondria-exclusion was testified by TOMM20 and β-actin immunoblotting (C).

(D-G) THP-1 cells were treated with Abt-S-Q and the transcription of IFN-β, IFN-λ, IL-6, TFN-α, and IFN-induced ADAR1p150 were quantified with RT-qPCR at 0, 8 and 16 h. In the mitochondria-excluded cytoplasm (E) the mtRNA (F) and mtDNA (G) were quantified.

(H and I) In Bax-knockdown A549 cells, Abt-S-Q-induced cytoplasmic mtRNA (H) and mtDNA (I) were quantified. Data presented as mean ± SD. n = 3 biological replicates. One-way ANOVA and two-way ANOVA were used. *$p < 0.05$, ** $p < 0.01$, *** $p < 0.001$, **** $p < 0.0001$.

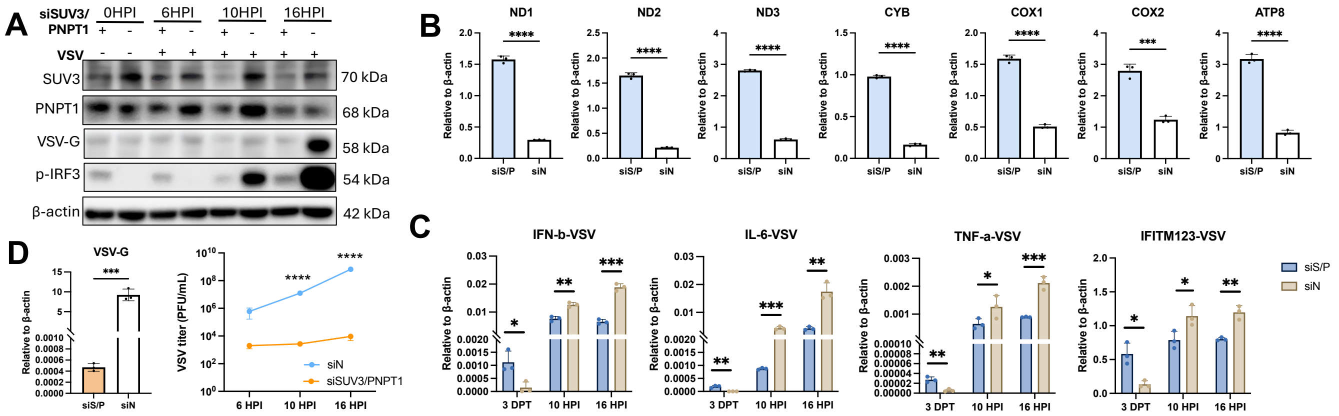


***Supplementary* Figure 2. The impaired mtRNA degradation results in mtRNA extrusion and restrained viral reproduction.**

(A-C) The SUV3- and PNPT1-knockdown A549 cells were infected with rVSV-GFP (MOI=0.005), the pIRF3 and viral protein G were detected by immunoblotting, the released mtRNA was quantified at 3-day post transfection (3 DPT), and the innate immunity-related transcripts were determined before and after rVSV-GFP-infection.

(D) The rVSV-GFP transcription was determined at 6 HPI and the infectious progeny virus was titrated by plaque assay at 6, 10 and 16 HPI. Data presented as mean ± SD. n = 3 biological replicates. Two-tailed Student's t test was used. *$p < 0.05$, ** $p < 0.01$, *** $p < 0.001$, **** $p < 0.0001$.

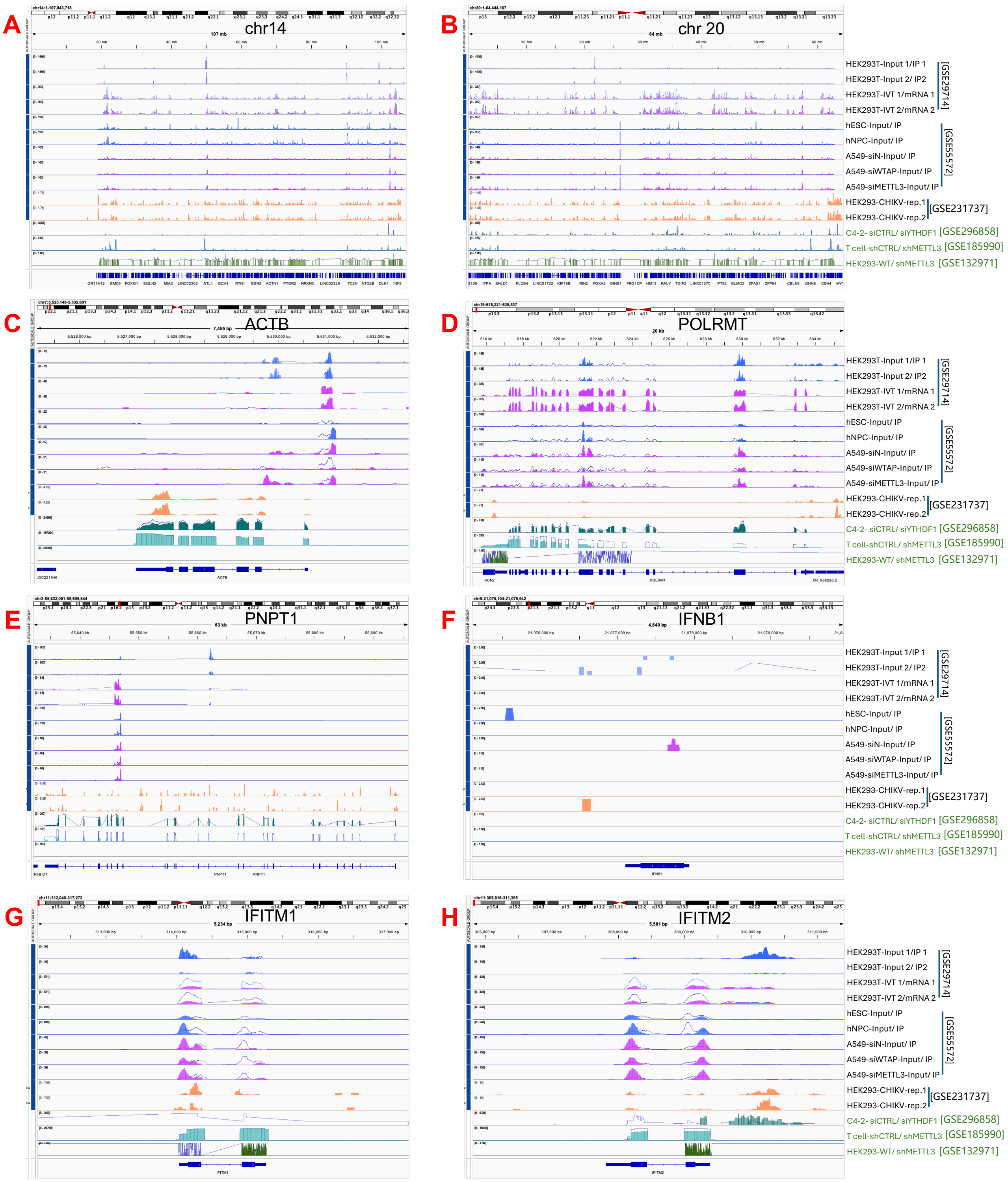


**_Supplementary_ Figure 3. The datasets from GEO revealed m$^6$A-modified nuclear-encoded transcripts.** (A-H) The m$^6$A modifications on the transcripts of Chromosome 14 (chr14, A) , Chr20 (B), ACTB (C), POLRMT (D), PNPT1 (E), IFNB1 (F), IFITM1 (G) and IFITM2 (H) were compared across multiple datasets from GEO database.

Table S1: siRNA oligonucleotide sequences, related to STAR methods.

| Genes (human) | Forward (5 '-3') | Reverse (5 '-3') |
|---|---|---|
| *RIG-I* | CGATTCCATCACTATCCAT | ATGGATAGTGATGGAATCG |
| *MDA5* | GTTATAGTTCTTGTCAATA | TATTGACAAGAACTATAAC |
| *TLR3* | GGTATAGCCAGCTAACTAG | CTAGTTAGCTGGCTATACC |
| *MyD88* | GCCTATCGCTGTTCTTGAA | TTCAAGAACAGCGATAGGC |
| *PKR* | GGTGAAGGTAGATCAAAGA | TCTTTGATCTACCTTCACC |
| *BAX* | GACGAACTGGACAGTAACA | TGTTACTGTCCAGTTCGTC |

Table S2: Primer sequence of target genes in qRT-PCR, related to STAR methods.

| Genes | Forward (5 '-3') | Reverse (5 '-3') |
|---|---|---|
| *IFNB*1 | CATTACCTGAAGGCCAAGGA | CAGCATCTGCTGGTTGAAGA |
| *IFNL*1 | CTAGACCAGCCCCTTCACAC | AAGGTGACAGATGCCTCCAG |
| *IL-6* | TACCCCCAGGAGAAGATTCC | TTTTCTGCCAGTGCCTCTTT |
| *TNF-a* | AACCTCCTCTCTGCCATCAA | CCAAAGTAGACCTGCCCAGA |
| *MAVS* | ATAAGTCOGAGGGCACCTTT | GTGACTACCAGCACCCCTGT |
| *IFITM123* | GACCTCCGTGCCCGACCATGT | CAGGGCCCAGATGTTCAGGC |
| *ADAR1p150* | CAATGAATCCGCGGCAG | GCCTTGAAATGGATGGGTGTAG |
| *ND1* | CCCATGGCCAACCTCCTACTCCTC | AGCCCGTAGGGGCCTACAACG |
| *ND2* | AACCCTCGTTCCACAGAAGCT | GGATTATGGATGCGGTTGCT |
| *ND3* | GACTACCACAACTCAACGGCTACA | ACTAAGAAGAATTTTATGGAGAAAGG |
| *ND4L* | CCCACTCCCTCTTAGCCAATATT | TAGGCCCACCGCTGCTT |
| *ND4* | ACAAGCTCCATCTGCCTACG | GCTTCAGGGGGTTTGGATGA |
| *ND5* | CTACCTAAAACTCACAGCCCTC | GGGTAGAATCCGAGTATGTTGG |
| *ND6* | GCCCCCGCACCAATAGGATCCTCCC | CCTGAGGCATGGGGGTCAGGGGT |
| *COX1* | GCCATAACCCAATACCAAACG | TTGAGGTTGCGGTCTGTTAG |
| *COX2* | ACCAGGCGACCTGCGACTCCT | ACCCCCGGTCGTGTAGCGGT |
| *COX3* | CCTTTTACCACTCCAGCCTAG | CTCCTGATGCGAGTAATACGG |
| *CYB* | CCCACCCTCACACGATTCTTTA | TTGCTAGGGCTGCAATAATGAA |
| *ATP6* | TTATGAGCGGGCACAGTGATT | GAAGTGGGCTAGGGCATTTTT |
| *ATP8* | CCCCATACTCCTTACACTATTCC | CGTTCATTTTGGTTCTCAGGG |
| *POLRMT* | CATCACCTACACCCACAACG | GTGCACAGAGACGAAGGTCA |
| *TFAM* | CCGAGGTGGTTTTCATCTGT | TCCGCCCTATAAGCATCTTG |
| *TEFM* | CATCCCTGTACTGGGCCTTA | AAGCAATCGGAAAGCTTCAA |
| *TRMT10C* | ACATAGCAATGGGCTGGAAG | TCTCTGTGCAAAGCACCATC |

| | | |
|---|---|---|
| *ELAC2* | CTTCCCAACTTCCAGCAGAG | CGGGGCTTATGTTGACAAGT |
| *FASTKD3* | GATGGAAACCCTGCCTGACA | GATCAAGAACCACCAGGGCA |
| *PNPT1* | CAACAAGCTTCAGTGGCAAA | CATAGCACTGGGTGTTGGTG |
| *SUV3L1* | TGGTTACGCCGATACATCAA | ATTGTGCACACCATCTTGGA |
